# Multi-attribute Decision-making is Best Characterized by an Attribute-Wise Reinforcement Learning Model

**DOI:** 10.1101/234732

**Authors:** Shaoming Wang, Bob Rehder

## Abstract

Choice alternatives often consist of multiple attributes that vary in how successfully they predict reward. Some standard theoretical models assert that decision makers evaluate choices either by weighting those attribute optimally in light of previous experience (so-called *rational models*), or adopting heuristics that use attributes suboptimally but in a manner that yields reasonable performance at minimal cost (e.g., the *take-the-best heuristic*). However, these models ignore both the possibility that decision makers might learn to associate reward with whole stimuli (a particular combination of attributes) rather than individual attributes and the common finding that decisions can be overly influenced by recent experiences and exhibit cue competition effects. Participants completed a two-alternative choice task where each stimulus consisted of three binary attributes that were predictive of reward, albeit with different degrees of reliability. Their choices revealed that, rather than using only the “best” attribute, they made use of all attributes but in manner that reflected the classic cue competition effect known as *overshadowing*. The time needed to make decisions increased as the number of relevant attributes increased, suggesting that reward was associated with attributes rather than whole stimuli. Fitting a family of computational models formed by crossing attribute use (optimal vs. only the best), representation (attribute vs. whole stimuli), and recency (biased or not), revealed that models that performed better when they made use of all information, represented attributes, and incorporated recency effects and cue competition. We also discuss the need to incorporate selective attention and hypothesis-testing like processes to account for results with multiple-attribute stimuli.

## Introduction

Choice alternatives often consist of multiple attributes whose reliability at predicting desired outcomes varies. Although attributes of apples such as color, size, and texture may predict their sweetness, some attributes may be more predictive (e.g., red apples may be much sweeter than green ones) than others (small apples may be only marginally sweeter than large ones). One strand of research on human decisionmaking has asked how multiple attributes are processed and used to make choices on the basis of prior experience. One approach, referred to as *rational models*, assumes that attributes are weighted and combined optimally, that is, in a manner that maximizes utility (Dawes, 1979; Dawes & Corrigan, 1974; Lee & Cummins, 2004; Simon, 1956, 1976). However, cognitive resource limitations often make optimal decision-making difficult (Simon, 1990; Oh et al., 2016;). Alternative *heuristic models* such as the *take-the-best model* offer suboptimal solutions that consider some attributes while ignoring others (Gigerenzer & Goldstein, 1996; Gigerenzer & Todd, 1999). Numerous studies have been conducted to evaluate which of these approaches characterize human decision makers but have not yet yielded a clear answer (Bergert & Nosofsky, 2007; Bröder, 2000, 2003; Lee & Cummins, 2004; Newell & Shanks, 2003; Newell, Weston, & Shanks, 2003; Oh et al., 2016; Rieskamp & Otto, 2006).

As useful as these models have been for characterizing multi-attribute decisions, a number of investigators have pointed out that they incorporate assumptions about underlying psychological processes that are implausible (Bergert & Nosofsky, 2007; Bobadilla-Suarez & Love, 2017; Dougherty, Franco-Watkins, & Thomas, 2008; Shanks & Lagnado, 2000). In this work, we consider the potential implications the field of *reinforcement learning* (RL) has for traditional models of choice. An emphasis on learning is especially apt because one recurring finding is that, rather than applying one strategy unconditionally, decision makers’ choice of strategy is *adaptive*, that is, it is sensitive to the information and reward structure of the task (Bröder, 2000, 2003; Newell, 2005; Rieskamp & Hoffrage, 2008; Rieskamp & Otto, 2006). Here we embellish traditional models of choice in two ways on the basis of recent research from the field of reinforcement learning.

First, models of choice such as the rational and take-the-best models generally take an “integrate-then-compare” strategy (Kable & Glimcher, 2009; Rangel, Camerer, & Montague, 2008), in which choice alternatives are represented at the level of attributes, the attributes of the two choice alternatives on the same dimension are compared, and the results of those comparisons are integrated to derive a value for each alternative (and then a decision) (Hunt, Dolan, & Behrens, 2014; Lim, O’Doherty, & Rangel, 2013). However, decisions can instead be made by directly comparing choice alternatives (e.g., a big red apple vs. a small green one). This strategy is consistent with the traditional choice theories of utility maximization that suggest that utilities are calculated and compared at the level of the alternative (Dai & Busemeyer, 2014; Tversky & Kahneman, 1992; von Neumann & Morgenstern, 1944).

Indeed, the multi-cue classification literature has shown that classification decisions can be made on the basis of direct associations between stimulus configurations and responses (Bayley, Frascino, & Squire, 2005; Yin & Knowlton, 2006). This may occur even when correct classification is a function of cues considered independently (i.e., the learning of configurations is formally unnecessary) (Goldfarb, Chun, & Phelps, 2016; Poldrack et al., 2001). That literature assumes that the formation of such configurations occurs over time and is part of what classifiers learn from performing the task (Johansen & Palmeri, 2002). The classification tasks used in these studies have a structure that is similar to those used to study choice (e.g.,Bergert & Nosofsky, 2007; Lee & Cummins, 2004), in that that they required extensive training over many trials. Thus it is reasonable to postulate that choices might also be represented at the level of alternatives, perhaps after experience with the task (Farashahi, Rowe, Aslami, Lee, & Soltani, 2017).

A second insight from reinforcement learning concerns the specific mechanisms via which learning occurs. Models developed in the RL framework assume that learning is driven by prediction error, that is, the difference between predicted and received rewards (Rescorla & Wagner, 1972), an assumption that has two important implications. First, rather than weighting experiences optimally RL models compute a weighted average of the value of choice alternatives on the basis of received rewards in a manner that assigns greater weight to recent experiences. In fact, people’s decisions often exhibit *recency effects* in which proximal experiences are weighted more than distal ones (Barron & Erev, 2003; Hertwig, Barron, Weber, & Erev, 2004). Second, rather than learning cues independently error driven learning brings rise to *cue competition*, a phenomenon widely observed in both animal conditioning (Bouton, 1993; Kamin, 1969; Mackintosh, 1975; Pearce & Hall, 1980; Wagner, 1969; Wasserman, Franklin, & Hearst, 1974) and human learning (Baker, Mercier, Vallée-Tourangeau, Frank, & Pan, 1993; Gluck & Bower, 1988; Kruschke, 2001; Waldmann & Holyoak, 1992). For example, *overshadowing* arises when a more valid cue suppresses the learning of a less valid one (Busemeyer, Myung, Jae, McDaniel, 1993; Kruschke & Johansen, 1999). In fact, RL models have received substantial support from both behavioral and neurobiological studies (Bayer & Glimcher, 2005; Daw, O’Doherty, Dayan, Seymour, & Dolan, 2006; Schultz, Dayan, & Montague, 1997). Note that when combined with computational techniques that consider participants’ individual choices, RL models can also characterize some of the variability in those choices that arise due to particular trial orders (Daw, Gershman, Seymour, Dayan, & Dolan, 2011; Doll, Shohamy, & Daw, 2015; Niv, 2009; O’Doherty, Dayan, Friston, Critchley, & Dolan, 2003).

Nevertheless, RL models face challenges of their own. Such models traditionally associate reward with the choice alternatives presented on each trial (Gershman, 2015; Niv et al., 2015) and indeed such models have enjoyed success in modeling simple decision tasks in which those alternatives varied on single attribute (e.g., color). However, this approach becomes inefficient as the number of attributes increases, a phenomenon known as the *curse of dimensionality* (Sutton & Richard, 1998). Dimensionality is a curse because the number of needed stimulus representations grows exponentially with dimensionality (e.g., the number of configural representations is 4 (2^2^) for stimuli with two binary dimensions, 8 (2^3^) for those with three dimensions, etc.) (Bellman, 1957). Of course, the fact that traditional attribute-wise decision models avoid this exponential explosion in the number of to-be-learned states highlights the fact that they and RL have complementary strengths and weaknesses: The former represent choices as the integration of attributes but assume perfect learning whereas the latter predicts recency effects and cue competition but often posits an unrealistic number of stimulus representations. Accordingly, here we follow the lead of other researchers (e.g., Jones & Cañas, 2010; Niv et al., 2015) by considering variants of RL models that associate reward with the *cues* of multi-attribute choice alternatives rather than the alternatives themselves.

The current study assessed decision making in scenarios in which choice options have a number of attributes, a situation that arguably characterizes many if not most real-world decisions. We ask three questions. First, do decision makers make use of all attributes or only the best one? Second, are choices represented and evaluated alternative-wise or attribute-wise? Third, do choices exhibit traditional RL phenomena such as recency effects and cue competition?

Our multi-attribute decision task combined elements from Lee and Cummins’s (2004) task. There were three binary stimulus attributes. This number of attributes allows a comparison of models that differ in the number of attributes they consider (e.g., the rational vs. the take-the best model) while also resulting in a number of configural stimulus representations that is sufficiently modest (2^3^ = 8) that decision makers could conceivably learn them during the course of the experiment (and thus engage in an alternative-wise vs. an attribute-wise strategy). We associated with each attribute a target weight indicating how important that attribute was for predicting reward (Oh et al., 2016). Reward probabilities were derived through a linear combination of attributes with varying amount of evidence provided by one stimuli over the other (Yang & Shadlen, 2007). Because attributes predicted reward independently, optimal performance did not require that participants encode stimulus configurations. Note that the structure of the stimuli has some similarity to those used in the study of category learning and memory systems mentioned above, such as the weather prediction task in which attributes varied in their usefulness for classification, learning occurred gradually with experience, and reward feedback was probabilistic (Gluck, Shohamy, & Myers, 2002; Knowlton, Squire, & Gluck, 1994; Kruschke & Erickson, 1994; Kruschke & Johansen, 1999; Kruschke, 1992; Lagnado, Newell, Kahan, & Shanks, 2006; Oh et al., 2016; Poldrack et al., 2001).

To assess what participants learned about the attributes, we analyzed their choices and response times during a testing block in which no feedback was provided. To assess the dynamics of learning (e.g., the existence of recency effects and cue competition), we fit six computational models to participants’ trial-by-trial choice data. To foreshadow the main findings, we found that participants generally learned to use all attributes and the correct relative rank of those attributes. And, their response times increased as the number of discriminating attributes increased, supporting the claim that participants evaluated choices at the attribute-wise level. Finally, that their decisions were best characterized by an attribute-wise RL model implies an effect of recent reward histories and cue competition. Indeed, participants’ choices reflected overshadowing in which stronger cues resulted in less learning of a weaker cue.

## Method

### Design

There were two between-participants conditions: partial and full (see below). Participants were randomly assigned to condition subject to the constraint that an equal number of participants were assigned to each condition.

### Participants

60 participants (35 women; mean age 20.1 years) from New York University undergraduate research pool participated for course credit. Informed consent was obtained from participants in a manner approved by the University Committee on Activities Involving Human Subjects.

### Materials

The task stimuli were described as aliens whose bodies were composed of three binary attributes: head (triangular or rectangular); body (light or dark); and tail (big or small). See Fig. 1A for an example.

### Procedure

The task consisted of 6 training blocks and a testing block, each 36 trials long. Before the start of training participants were informed that all three attributes (head, body and tail) were predictive of reward and that they needed to learn about the importance of the cues and stimulus attributes through trial-and-error, with the goal to collect as many artificial one-dollar bills as possible. Participants were also informed about the probabilistic nature of the task. Specifically, they were told that “There was no perfect body part relating to reward, but you should choose the stimuli that you think is more likely to be rewarded.”

We first describe the full condition and then describe how it differed from the partial condition. During each training trial participants were presented with two schematic aliens (Fig. 1A). The participants’ task was to choose the alien that they deemed more likely to be rewarded. Participants had up to 5 s to make a response, after which the chosen stimulus was highlighted by a red frame for 1 s. The reward associated with that choice was displayed in the center of the screen and consisted of an image of either a one-dollar bill or zero-dollar bill. The stimuli and the reward remained on the screen for 3 s, after which a fixation cross was displayed on a blank screen for 3 s (Fig. 1B). There were six possible mappings from physical (head, body, and tail) to logical attributes (Attribute 1, 2 and 3). These mappings were counterbalanced such that the same number of participants was assigned to each of the six mappings.

**Fig. 1.**
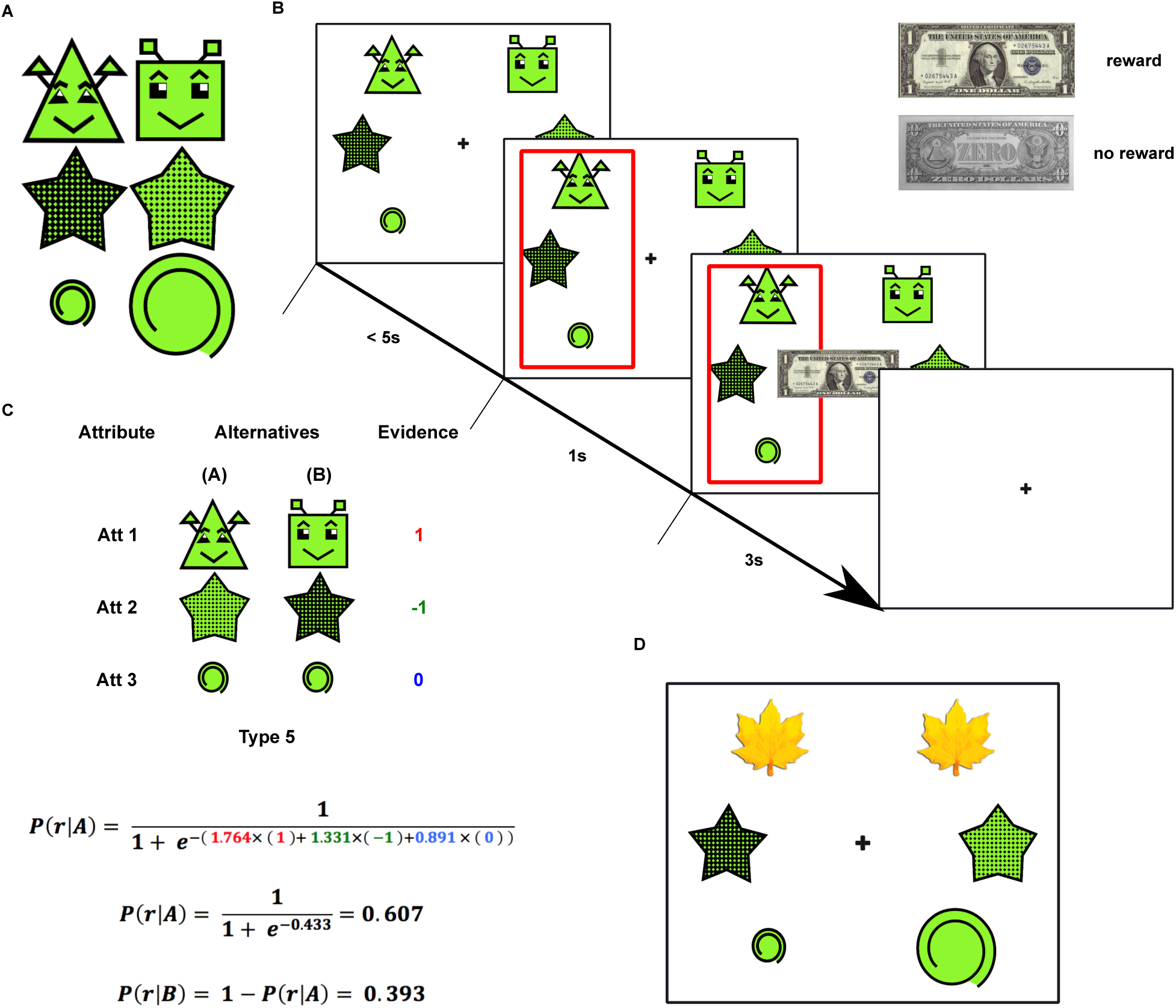
Methods overview. (A) Stimuli consisted of features that comprised each of three binary dimensions of alien body parts: head (triangular or rectangular); body (light or dark); and tail (big or small). (B) Schematic of a single trial. On each trial, participants were presented with 2 stimuli, each having a feature along each of the three body parts. Participants then chose one of the stimuli and received binary outcome feedback, shown as one-dollar bill (reward) or zero-dollar bill (no reward). The next trial began after a delay. (C) An example of reward probability computation. Reward probabilities were derived through a logistic regression, where evidence of attributes and corresponding attribute weights were linearly combined. (D) Illustration of a trial in the partial condition. On this trial, the two heads are covered by pieces of leaves.

To determine reward on each trial, we associated with each stimulus attribute a target weight indicating how important that attribute was for predicting reward. Those weights were 1.778, 1.333, and 0.889, for Attributes 1, 2 and 3, respectively. These attribute weights are compensatory such that the two least valid Attributes 2 and 3 together outweighed the most valid Attribute 1. We defined 13 types of choice problems defined by the amount of evidence provided by one alternative (referred to as A) over the other (B). For each choice type, Table 1 defines whether the evidence provided by the cues on Attribute *i,* which we will refer to as *Ev*(*Att_i_*), favor alternative A or B. When A and B display the same cue on an attribute then of course it favors neither alternative and so *Ev*(*Att_i_*) = 0. But when the two cues differ, *Ev*(*Att_i_*) = 1 means that alternative A displays the cue more predictive of reward whereas *Ev*(*Att_i_*) = −1 means that B does. Some choice types could be instantiated in multiple ways. For example, choice type 1 (characterized by *Ev*(*Att*_1_) = 1 and *Ev*(*Att*_2_) = *Ev*(*Att*_3_) = 0) could be instantiated in four ways: {aaa, baa}, {aab, bab}, {aba, bba}, or {abb, bbb}, where each set denotes alternatives A and B and each “a” and “b” are the cues that favors A and B, respectively. Choice types 1-3 had four instantiations, types 4-9 had two, and types 10-13 had one.

**Table 1.** Structure of the task.

| Choice Type | $Ev(Att_1)$ | $Ev(Att_2)$ | $Ev(Att_3)$ | Target<br>Pr ($r\vert A$) | N | Rewarded | Actual<br>Pr ($r\vert A$) |
| --- | --- | --- | --- | --- | --- | --- | --- |
| 1 | 1 | 0 | 0 | 0.855 | 24 | 21 | 0.875 |
| 2 | 0 | 1 | 0 | 0.791 | 24 | 19 | 0.792 |
| 3 | 0 | 0 | 1 | 0.709 | 24 | 17 | 0.708 |
| 4 | 1 | 1 | 0 | 0.957 | 12 | 11 | 0.917 |
| 5 | 1 | -1 | 0 | 0.609 | 12 | 7 | 0.583 |
| 6 | 1 | 0 | 1 | 0.935 | 12 | 11 | 0.917 |
| 7 | 1 | 0 | -1 | 0.709 | 12 | 9 | 0.750 |
| 8 | 0 | 1 | 1 | 0.902 | 12 | 11 | 0.917 |
| 9 | 0 | 1 | -1 | 0.609 | 12 | 7 | 0.583 |
| 10 | 1 | 1 | 1 | 0.982 | 18 | 18 | 1.000 |
| 11 | 1 | 1 | -1 | 0.902 | 18 | 16 | 0.889 |
| 12 | 1 | -1 | 1 | 0.791 | 18 | 14 | 0.778 |
| 13 | -1 | 1 | 1 | 0.609 | 18 | 11 | 0.611 |

Note. 13 choice types were constructed. For each attribute, evidence score ( $E_i$ ) (columns 2-4) indicate if the cues on Attribute $i$ favor alternative A ( $Ev(Att_i) = 1$ ), alternative B ( $Ev(Att_i) = -1$ ), or do not discriminate between alternatives ( $Ev(Att_i) = 0$ ).

Target reward probabilities were derived from a logistic regression in which the evidence provided by each attribute was linearly combined (Yang & Shadlen, 2007):

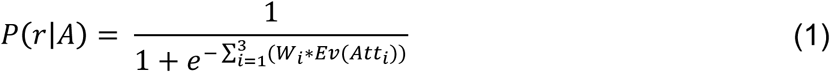

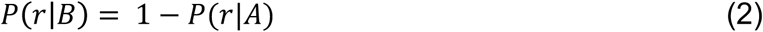

where *P*(*r*|*A*) represents the probability of reward given that stimulus A is chosen. The target *P*(*r*|*A*) for each of the 13 choice types is presented in Table 13. It also presents the number of times each choice type was presented during the 216 training trials (e.g., choice type 1 was presented 24 times). The number of times each choice type was rewarded was chosen so as to approximate its target reward probability as closely as possible. For example, to approximate its target reward probability of .855, choice type 1 was rewarded on 21 of its 24 presentations. Table 1 also presents the actual probability of reward for each choice type (e.g., actual *P*(*r*|*A*) = 21 / 24 = .875 for choice type 1). The attribute weights recoverable from these 216 training trials were 1.764, 1.331 and 0.891. The compensatory feature of these weights is illustrated by choice type 13, in which the cues on Attribute 1 implicate B as the best choice whereas those on Attributes 2 and 3 implicate A. Because *W*_2_(1.331) + *W*_3_(0.891) > *w*_1_(1,764), an “A” response was more likely to be rewarded (11 / 18 = 0.611). As will become clear later, choice type 13 will serve as a direct behavioral measure of overshadowing.

Fig. 1C presents how reward probabilities were determined for choice type 5. In this example head, body and tail correspond to Attribute 1, 2 and 3, respectively, and triangular head, dark body and small tail are the cues that favor alternative A over B. Whereas the cues on Attribute 1 favor A (*Ev*(*Att*_1_,) = 1, red), those on Attribute 2 favor B (*Ev*(*Att*_2_) = −1, green), and those on Attribute 3 do not discriminate between alternatives (*Ev*(*Att*_3_) = 0, blue). These sources of evidence were then linearly combined with their corresponding attribute weights to derive reward probabilities (Eqs. 1 and 2 in the bottom panel of Fig. 1C).

Note that it is useful to recode the attributes weights of 1.764, 1.331 and 0.891 as a vector of normalized weights *W^n^* (.443, .334, and .224) and a scaling (or “inverse temperature) parameter *β^n^* (4.0). The normalized weights reflect the relative importance of the three attributes that should be adopted by any ideal observer. Of course, maximizing reward entails that participants should always select the choice alternative that is more likely to be rewarded (i.e., their decision rule should use *W^n^* but adopt a scaling parameter *β* = ∞). But as in many studies of choice we expect decision makers to also display a degree of decision noise. Logistic regression analyses of our participants’ choices will yield attribute weights that can also be recoded as normalized weights and a scaling parameter *β*. The normalized weights will reveal how many attributes participants use to make choices, whether their attribute use reflects the objective validity of the attributes (Attribute 1 > 2 > 3), and more subtle features of their weights such as overshadowing (a relatively higher weight on the high-validity Attribute 1 and a relatively lower one on the low-validity Attribute 3). Their scaling parameter will be interpreted as reflecting decision noise, where a smaller value (or higher temperature) of *β* reflects greater noisier decisions (i.e., more probability matching) and a larger one (or lower temperature) reflects more deterministic responding.

Instantiations of the 13 choice types were assigned to blocks of 36 training items such that the number of each choice type was the same in each block. There were 4 instances each of choice types 1–3 in each block, 2 instances each of choice types 4–9, and 3 instances each of choice types 10–13.

The partial condition was identical to the full condition except that on some trials an attribute in both alternatives was covered and so unobservable. For instance, on a given trial, participants might be shown two aliens, with both of their heads covered by pieces of leaves (see Fig. 1D). Because we found that the full versus partial manipulation yielded few differences, details of the stimuli in this condition are presented in Appendix A.

The testing phase of the experiment consisted of a single block of 36 trials. All participants could see all cues on all attributes. The procedure was identical except that feedback was omitted.

### Computational models

We constructed six computational models that differed in three aspects: attribute use (best attribute only vs. all attributes), choice representation (alternative-wise vs. attribute-wise), and whether the models reflect error driven learning. We first present the three models that assume perfect learning and then three that incorporate recency effects and cue competition.

The *attribute-wise, full-learning model* (Att-FL) is a variant of a weighted additive, rational decision model, *RAT* (Bergert & Nosofsky, 2007; Dawes & Corrigan, 1974; Lee & Cummins, 2004; Rieskamp & Otto, 2006). It uses all the relevant available information to update cue validities on the basis of the proportion of rewarded inferences made across stimulus pairs in cases where a cue discriminates between alternatives. The validity of cue *i* on trial *t* was defined as:

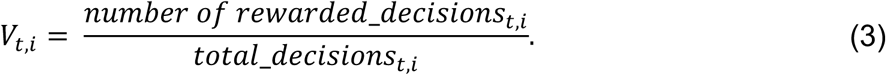

A problem with this definition identified by Lee and Cummins is that the validity of a cue that makes 1 out of 1 rewarded decisions equals that of a cue that makes 100 out of 100 rewarded decisions. Lee and Cummins addressed this problem with the Bayesian modification given in Equation 4:

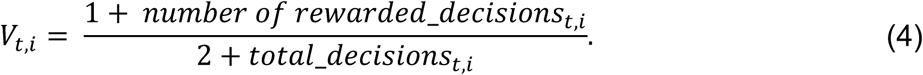

Thus, cues that make a smaller number of rewarded decisions have a lower Bayesian cue validity. We adopted this definition of cue validity in the current study. According to Equation 4, on each trial the total number of decisions made by every discriminating cue that appeared in both alternatives is incremented by 1. In addition, if the chosen stimulus was rewarded, the number of rewarded decisions for each of its cues was increased by 1. Because participants were instructed that only one stimulus was rewarded on each trial, the number of rewarded decisions for each cue in the unchosen stimulus was increased by 1 when the chosen stimulus was not rewarded.

Cue *weights* were then calculated as the log odds of the cue validities, representing each cue’s independent contribution in favor of an alternative (Katsikopoulos & Martignon, 2006; Lee & Cummins, 2004) in Equation 4.

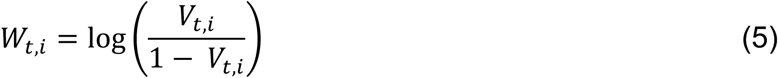

The value of each stimulus *j* was determined by the sum of cue weights for that stimulus:

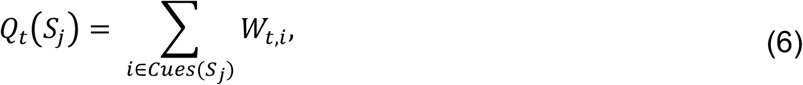

where *Cues(S_j_)* denotes the cues on *S_j_*.

The original RAT model defined by Lee and Cummins (2004) assumed a deterministic decision rule in which a stimulus was always chosen when its *Q* value exceeded that of the other stimulus. Here we follow Bergert and Nosofsky (2007) by defining a “noisy” version of RAT in which the probability of choosing one stimulus increases as the degree of evidence in favor of that stimulus increases. In particular, the *Q* values were entered into the softmax choice function

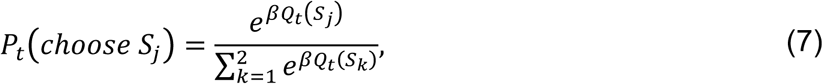

where the inverse temperature parameter *β* again represents the level of noise in the decision process, with larger values of *β* corresponding to low decision noise and near-deterministic choices and smaller ones corresponding to high decision noise and nearly random decisions. This model can be thought of an “ideal observer” learning model (albeit one with decision noise) that optimally evaluates the probability of an alternative getting rewarded given cues and associated validities for both alternatives. In particular, this model assumes perfect learning and uses all the attributes to evaluate choice at the level of attributes.

The *take-the-best, full learning model* (TTB-FL) also assumes perfect learning, using the same rule to update cue validities as Att-FL represented by Equations 4 and 5. However, instead of deciding on the basis of all cues, TTB-FL sequentially searches through cues in descending order of their validities and stops upon reaching a cue that discriminates the alternatives (Gigerenzer & Todd, 1999). The weights for that cue and the cue in the other alternative on the same attribute are taken as the value for those alternatives and entered in the softmax function above to compute choice probabilities.

Although it also assumes perfect learning, *the alternative-wise, full learning model* (Alt-FL) differs from the models above in how it represents and updates stimulus values. Instead of treating each alternative as consisting of three attributes, it treats it as a whole stimulus and computes the validity of stimulus in a manner analogous to the Att-FL model.

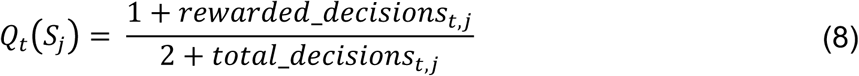

The *Q* values are then entered the softmax function. When a reward was received, the number of rewarded decisions of the chosen stimulus was increased by 1. When no reward was received, the number of rewarded decisions of the unchosen stimulus was increased instead. The total number of decisions of both stimuli were increased by 1 on every trial. Note that in the present task the total numbers of distinct stimuli presented are 8 and 20 in the full and partial conditions, respectively.

We now describe models that incorporate error driven learning. The *attribute-wise, RL model* (Att-RL) assumes that the value of a stimulus S presented on trial *t* is calculated as the sum of the values of its cues on that trial. For consistency of notation with the previous models, we use *W* to refer to a cue’s value or “weight”.

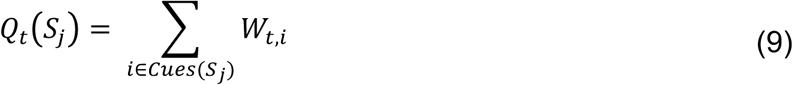

The *Q* values are then entered the softmax function.

After feedback is received, the weights of the cues are updated according to the standard Rescorla-Wagner learning rule (Rescorla & Wagner, 1972). The prediction error *δ* on trial *t* was calculated as the difference between the reward expected on the basis of the chosen stimulus and the reward received:

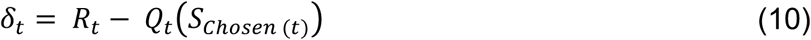

where *S_chosen(t)_* is the stimulus chosen on trial t. *δ=* was then used to update the weight of each cue *i:*

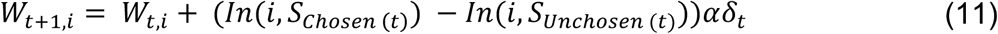

where *α* is a learning-rate parameter and *In(i,S_j_)* returns 1 when *i ∈ Cues(Sj)* and 0 otherwise. Note that when the prediction error *δ_t_* is positive then the weights of the cues in the chosen stimulus increase and those in the unchosen stimulus decrease. When *δ=* is negative the weights change in the opposite direction. A cue’s weight is left unchanged if it appears in both or neither stimuli. Eq. 11 updates the cues in a manner that recent experiences are weighed more heavily than distal ones. Later we demonstrate that Att-RL predicts the competition among cues that results in overshadowing.

The *take-the-best, recency-weighted learning model* (TTB-RL) assumes that cue weights are learned as in the Att-RL model (Eq. 11). However, it uses the rule defined by TTB-FL to make decisions. That is, only the weight of highest-ranked cue (and the other cue in the same attribute) determines the *Q* values for the two alternatives when cues discriminate the alternatives.

Finally, the *alternative-wise, recency-weighted learning model* (Alt-RL) learns values for each stimulus following the R-W rule. On each trial, the values of each stimulus was updated according to:

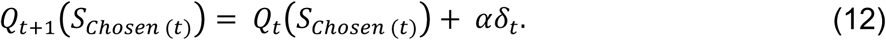

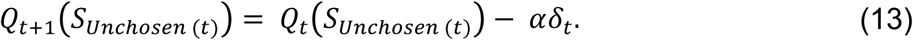

such that the values of the chosen and unchosen stimuli increase and decrease, respectively, when *δ_t_* is positive and vice versa when *δ_t_* is negative.

Together, these six models allow a quantitative assessment of how multi-attribute decisions are represented, how attributes are used, and whether decisions are overly influenced by recent experiences (Table 2).

**Table 2.**
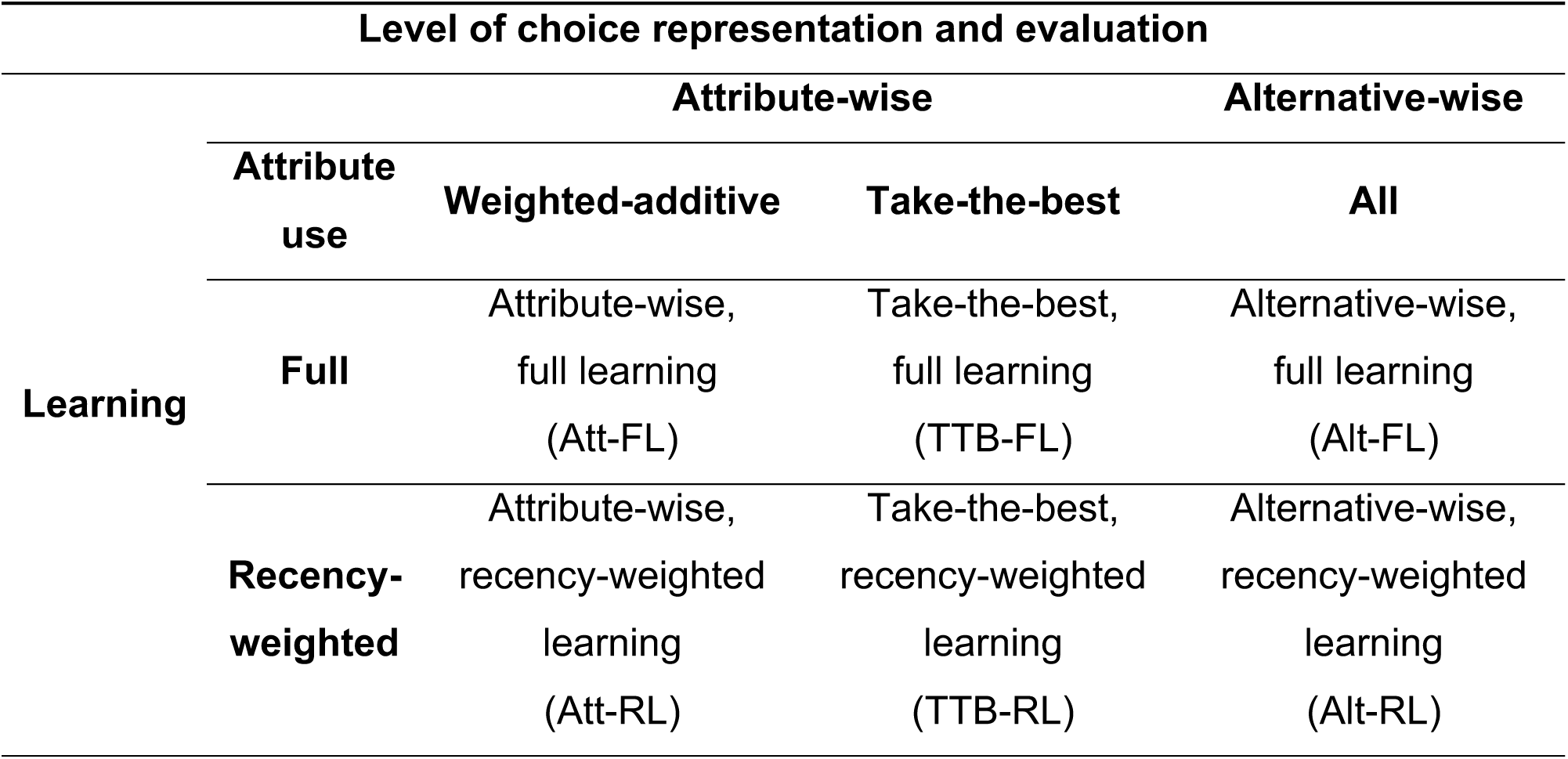
Computational models.

## Results

### Learning

To examine learning, we defined an optimal choice as one that maximizes reward over the experiment (see Table 1). Participants were excluded if their percentage of optimal choices failed to reach a criterion of 60% in the last 2 blocks of the experiment, which suggested they failed to learn the task. 13 participants were excluded, leaving a total of 47 participants for further analysis.

Participants’ performance improved during the experiment. The mean proportions of optimal choices for each block of 36 trials are shown in Fig. 2. An ANOVA with block as a within-participant factor and learning condition (complete vs. partial) as a between-participant factor revealed a main effect of block, *F*(6, 270) = 12, *MSE* = 0.007, p < 10^−7^, no effect of condition, *F*(1, 45) = 0.03, *MSE* = 0.039, *ns,* and no effect of a Block X Condition interaction, F(6, 270) = 0.51, *MSE* = 0.007, *ns.* Thus, we collapse the full and partial conditions in analyses that follow.

**Fig. 2.**
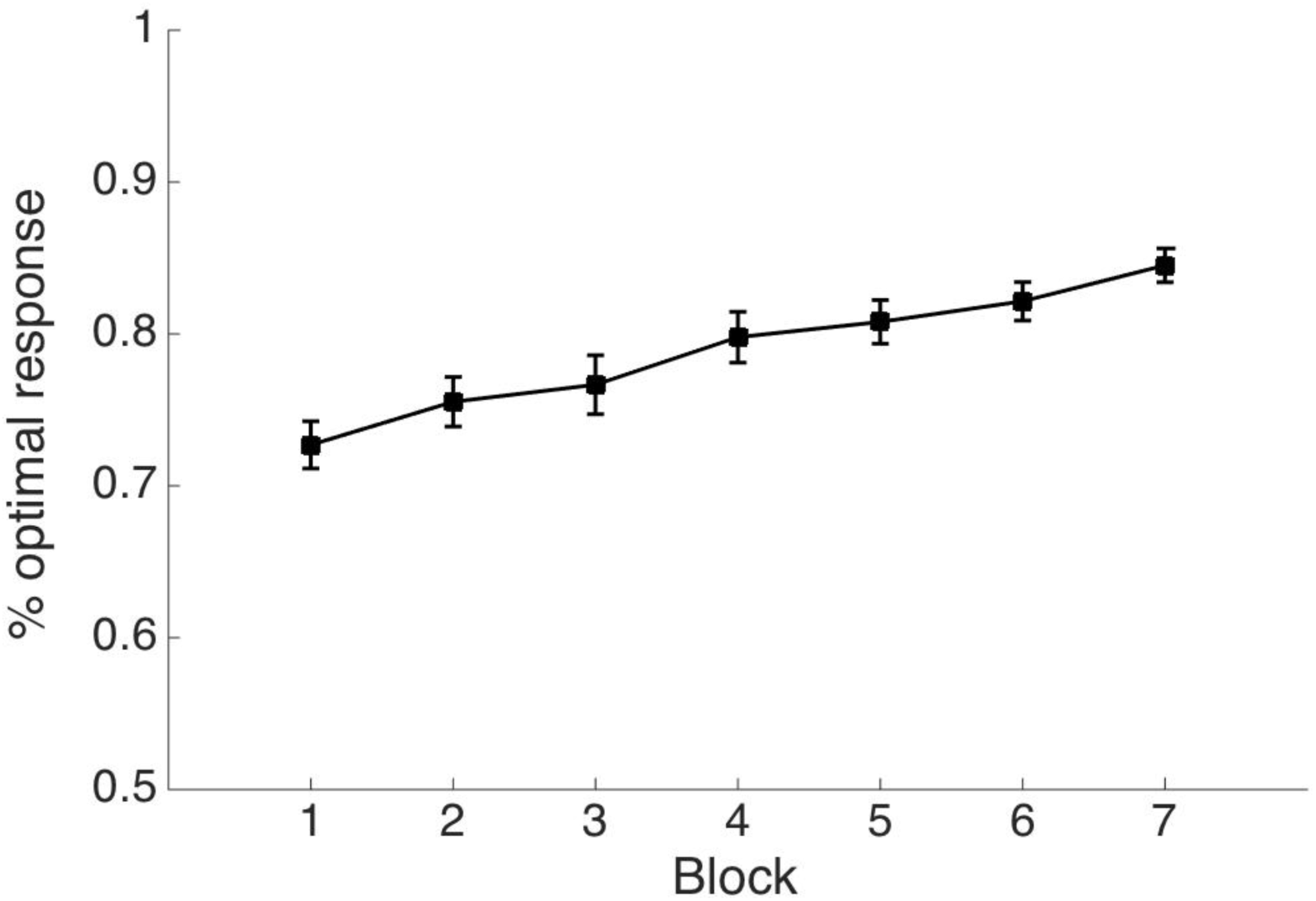
Learning across blocks and learned participants. Plotted is the proportion of optimal responses. Error bars represent ± 1 sEm across participants.

### Attribute Use During Test

Although a central goal of this article is to evaluate alternative learning models of choice, this section aims to characterize how participants used attributes to make choices after six block of training without regard to how those attributes were learned. To this end, we first fit a *linear weighted additive model*, or *WADD* (Payne, J. W. Bettman, J. R. Johnson, 1993), in which the following logistic regression was used to derive subjects’ attribute weights *W_i_* on the basis of evidence provided in each choice type (Table 1). ^1^

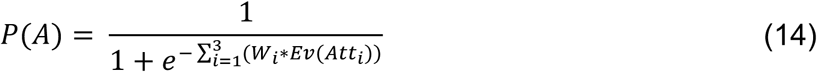

Estimating the attribute weights involved using a variational Bayesian method (Drugowitsch, 2013). This method was chosen to solve complete separation problems in traditional logistic regression analysis with small sample sizes (Gelman, Jakulin, Pittau, & Su, 2008), by assigning a hyper-prior to each individual regression weight. In our analysis, the hyper-prior was specified as Gamma (10^−2^, 10^−4^), such that the prior of regression weights was not informative (Drugowitsch, 2013).

The weights averaged over participants were 3.339, 2.128, and 1.358 for Attributes 1, 2, and 3, respectively. These weights reflect two important findings. The first is that after six blocks of training all three attributes were influencing participants’ choices. The second is that learners also recovered the attributes’ relative ranking. The weight on Attribute 1 was statistically greater than that on Attribute 2, t(46) = 5.557, p < 10^−5^, which in turn was greater than that on Attribute 3, t(46) = 3.880, p < 10^−3^, which in turn was greater than 0, t(46) = 6.103, p < 10^−6^.

Although these results characterize group level performance, it is important to ask if they describe most participants’ performance or are a result of averaging over participants with very different performance profiles. To this end, each learner’s attribute weights were normalized and the results are displayed in the simplex plot in Fig. 3A (Coenen, Rehder, & Gureckis, 2015). Points within the simplex reflect the relative contribution of the three attributes on participants’ choices. The middle asterisk (black) corresponds to the case where three attributes have equal influence on decisions, the red asterisk depicts the optimal normalized weights (0.443, 0.334 and 0.224) and the magenta asterisk shows participants’ average normalized weights. Informal inspection of the distribution of simplex plot points reveals that participants generally recovered the attributes’ relative importance. Of the six possible orderings of attribute weights, the weights of 26 out of 47 (55%) participants reflected the optimal ordering (Attribute 1 > 2 > 3; Fig. 3B). Furthermore, Attributes 1, 2 and 3 were the most heavily weighted attribute for 85%, 11% and 4% of the participants, respectively.

**Fig. 3.**
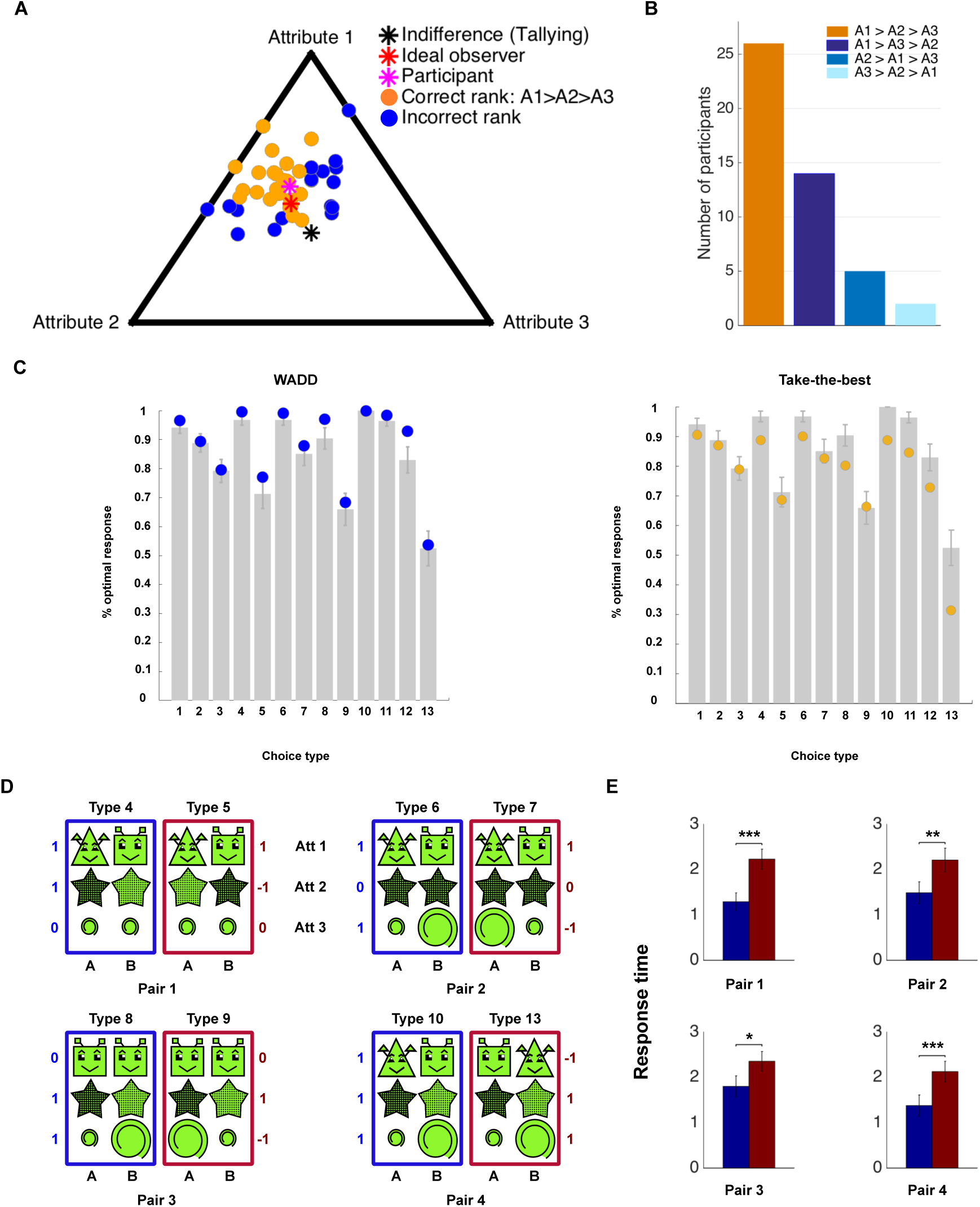
Attribute use. (A) Simplex plot of normalized attribute weights. (B) Histogram of the number of participants with different ranks of attributes. (C) Examples of choice pairs used to contrast response time predictions of the take-the-best and rational models. In these examples the head, body, and tail correspond to Attributes 1, 2, and 3, respectively. The amount of evidence provided by alternative A over B is presented alongside the three attributes. *Ev*(*Att_i_*) = 0 when A and B display the same cue on an attribute, 1 when alternative A displays the cue more predictive of reward, and −1 when B does. (D) RTs for the four choice pairs shown in (C).

That participants placed a substantial weight on all three attributes provides preliminary evidence against the take-the-best model, which predicts that most decisions are determined by the stronger cues. To formally evaluate the take-the-best model as an account of participants’ decision strategy, we also fit it to their test block choices. To give this strategy additional flexibility (and allow a more direct comparison with WADD), we followed the lead of Bergert and Nosofsky (2007) and fit a version of take-the-best in which attributes are not assumed to be learned perfectly (as in the standard take-the-best model) but rather are free parameters. This model thus has four free parameters (three attribute weights and a scaling parameter for the softmax choice rule; see Eq. 7). The predictions of both this model and WADD are shown in Fig. 3C (yellow and blue plot points, respectively) superimposed on the empirical data (gray bars). The figure reveals that even with fitted attribute weights, take the best is a poor account of those choice types that can be influenced by the cues on multiple attributes. For example, choice types 4, 6, 8, and 10-12 are all examples of choices in which the cues on multiple attributes favor of alternative A. But because take-the-best decides on the basis of only one attribute, it systematically underestimates the choice probabilities of those choice types. It also favors choice alternative B on the choice type 13 (because B is implicated by the strongest Attribute 1) whereas participants’ choices reflected indifference (~0.5). In contrast, WADD provided a superior account of each of these choice types. As a result, the average log likelihood of the take-the-best model was lower than that of WADD (−11.129 vs. −7.736), despite having an extra parameter. Remarkably, WADD was the better fitting model for every one of the 47 participants. Another version of take the best with a free weight parameter for each of the six cues fared no better.^2^

Bergert and Nosofsky (2007) considered yet another generalization of take-the-best, which was to assume that the single attribute whose cues are initially compared is not always the “best” attribute but rather is chosen probabilistically in a manner that reflects those weights, so that the best attribute is chosen with highest probability, the second-best is chosen with the second highest probability, and so forth. Because Bergert and Nosofsky observed that the choice predictions of this generalization of take-the-best can be indistinguishable from a model that chooses on the basis of weighted attributes (like WADD), they conducted a novel response-time (RT) analysis. This analysis makes use of the well-known finding that more difficult choices take longer to make (Gold & Shadlen, 2007; Hunt et al., 2014). We identified pairs of choice types in which the evidence for one alternative over another was identical if only the “best” attribute was considered. For example, although on the most valid Attribute 1 choice types 4 and 5 both favor Alternative A, on Attribute 2 choice type 4 favors A whereas 5 favors B (Table 1). Because take-the-best only considers the best discriminating attribute, it predicts that choice types 4 and 5 are equally difficult and so made in the same amount of time. In contrast, models that consider all attributes predict that choice type 4 is easier than (and so made faster than) type 5. Bergert and Nosofsky referred to choice types like 4 and 5 as *RAT-easy* and *RAT-hard* problems, respectively (because RAT is an example of a model that makes use of all attributes). Other pairs of RAT-easy and -hard choice types are 6 and 7, 8 and 9, and 10 and 13 (Fig. 3D). In fact, the RAT-easy choices (4, 6, 8, and 10) required less time on average than the corresponding RAT-hard ones, t(46) = −4.61, p < 10^−3^; t(46) = −3.48, p < 0.005; t(46) = −2.54, p < 0.05; t(46) = −4.50, p < 10^−3^, respectively. Figure 3E presents the RTs for all four pairs. These RT analyses corroborate the conclusion drawn above on the basis of the choice data, namely, that participants were not “taking the best” after six blocks of training.

That participants apparently made use of all three attributes in an added weighted fashion led us to additionally ask how those attribute weights compared to those of an ideal observer. Participants’ average normalized attribute weights derived from WADD model (Eq. 14) were 0.503, 0.309, and 0.188 (Fig. 3A, the magenta asterisk) as compared to the optimal normalized weights of 0.443, 0.334 and 0.224 (Fig. 3A, the red asterisk). That is, they overweighed Attribute 1 and underweighted Attribute 3. Recall that this pattern of attribute use is consistent with the cue competition effect known as overshadowing in which the presence of strong cues (e.g., those on Attribute 1) result in reduced learning of weaker cues (e.g., those on Attribute 3). To assess the presence of overshadowing statistically, we computed the linear trend in each subject’s attribute weights by subtracting the weight for Attribute 3 from that of Attribute 1. We then compared that linear trend against the linear trend in the normalized weights: 0.443 − 0.224 = 0.219. The result—1(46) = 4.461, p < 10^-4^—supports the conclusion that Attribute 1 was overweighed and Attribute 3 was underweighted relative to the ideal weights.

Although we attribute this pattern of attribute weights to error driven learning, it important to ask whether it resulted from a decision strategy instead. Oh et al. (2016) found that in order to cope with time pressure participants adopted a strategy in which they dropped less valid attributes (also see Lee & Cummins, 2004). Because we imposed a 5 s response deadline, it is conceivable that time pressure reduced our participants’ relative use of Attribute 3 and so increased their relative use of Attribute 1. To assess this possibility, we examined response times during the test block. Participants took an average of 1.815 s (SD = 0.980) to respond; the RT for the slowest choice type 9 was 2.355 (SD = 1.145) s. That our participants responded well before the response deadline supports the conclusion that the presence of overshadowing was not the result of a decision strategy induced by time pressure.

To demonstrate the effect of overshadowing on participants’ choices during the test block, we predicted those choices with a variational Bayesian logistic regression model in which the attribute weights were stipulated to be the normalized ideal weights (i.e., 0.443, 0.334 and 0.224) but included a single free scaling parameter *β*, that is,

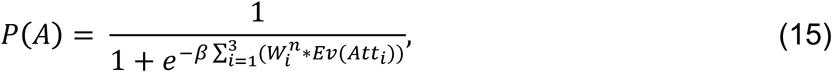

where 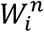 are the normalized ideal weights. The top left panel of Fig. 4B presents the probability of responding optimally as predicted by this ideal observer model (red circles) superimposed on participants’ actual performance (gray bars). Although this model predicts participants’ choices fairly well, it mis-predicts the choice type that serves as a direct test of compensatory weights, choice type 13. Whereas the ideal observer model predicts that alternative A in this choice should be favored, participants’ responses did not differ significantly from 0.50, M = 0.525, t(46) = 0.415, ns (Fig. 4B). This misprediction arises of course because whereas the optimal weights are compensatory (Attribute 1 ≈ Attribute 2 + Attribute 3; Fig. 4A, upper panel), participants’ weights derived from WADD were not (Attribute 1 « Attribute 2 + Attribute 3; Fig. 4A, lower panel). Indeed, a paired-t test conducted on those weights revealed no statistical difference between Attribute 1 and the sum of Attributes 2 and 3, t(46) = 0.266, ns. Comparison of the fit of the ideal observer model to that of WADD (blue circles in Fig. 4B) confirms that the latter’s non-compensatory weights reproduces the ~0.5 choice probability on choice type 13.^3^

**Fig. 4.**
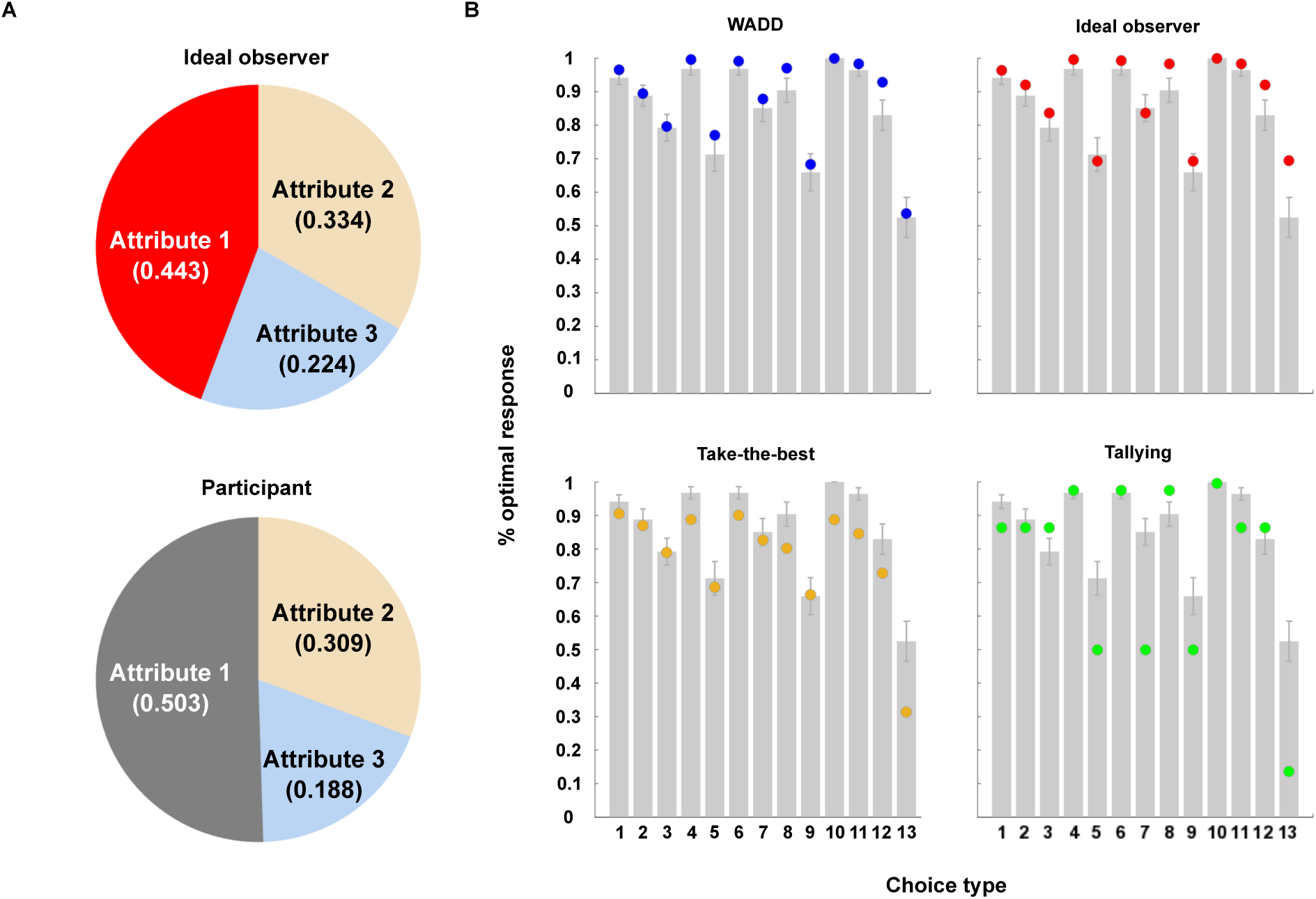
Attribute weights. (A) Normalized attribute weights inferred from an ideal observer model and those of participants. Ideal observer model predicts compensatory weights whereas participants’ weights were non-compensatory. (B) Participants’ performance and predictions from four decision models: three-parameter full model (Eq 13), the ideal observer model (Eq. 14), the take-the-best model and the tallying model.

For completeness, Fig. 4B also presents the predictions of another heuristic known as *tallying* (Dawes, 1979; Gigerenzer & Gaissmaier, 2011). According to tallying, one simply counts the number of attributes favoring one alternative over the other (Gigerenzer & Gaissmaier, 2011). (For this reason, tallying is sometimes referred to as an *equal weight heuristic;* (Bröder, 2000; Dawes, 1979; Payne, J. W. Bettman, J. R. Johnson, 1993) To make its predictions comparable to the other models, we granted tallying a free scaling parameter *β*. Unsurprisingly given our result regarding relative attribute use, tallying (green circles in Fig. 4B) was also a poor account of participants’ choices (e.g., it predicts chance performance on choice types 5, 7, and 9, that alternative B should be favored for choice type 13, etc.).

Finally, recall that the unnormalized attribute weights that emerge from the logistic regression analysis in Eq. 14 can be recoded so as to yield not only normalized weights but also a scaling parameter *β*. Participants’ average value of *β* was 5.756, which is larger than that used to generate the training data (4). (In the single parameter ideal observer model of Eq. 15, the average best fitting value of *β* was 7.339). That is, participants’ choices reflected responding that was more decisive than that implied by pure probability matching (Estes, 1976; Lagnado et al., 2006; Vulkan & Evolution, 2000).

In summary, participants learned that all three attributes were predictive of reward and the relative rank of those attributes. They overweighed the most predictive attribute (Attribute 1) and underweighed the least predictive one (Attribute 3), a fact that is consistent with overshadowing and that resulted in learned weights that were not compensatory and at-chance performance on choice type 13. Yet, if one ignores choice type 13, the choices of a large majority of the participants were not dramatically different than those implied by the ideal attribute weights. And, their choices reflected a relatively low level of probability matching. Overall, participants learned the task reasonably well.

### Attribute-wise versus Alternative-wise Representations

Although the preceding analyses indicate that participants made use of all information, it doesn’t directly address whether choice alternatives were represented at the level of attributes or alternatives. To answer this question, we carried out an additional RT analysis. Because previous work in the probabilistic classification literature suggests that use of whole-stimulus representations emerged with task experience (Gluck et al., 2002; Johansen & Palmeri, 2002; Poldrack et al., 2001), we again restrict our analysis to the testing block. Recall that the 13 choice types differed in number of attributes that discriminated the alternatives (i.e., number of attributes for which *Ev(Att_i_*) Ψ 0). The alternatives in choice types 1-3 differ in one attribute, 4-9 differ in two, and 10-13 differ in all three. Attribute-wise models predict longer RTs for choice types with more attributes that discriminate, all else being equal (see below). In contrast, alternative-wise models predict that RTs should not vary with the number of discriminating attributes. This is so because they stipulate that integrated values of whole stimuli are compared.

However, a simple analysis in which RTs are predicted from the number of discriminating attributes (hereafter referred to as *nAtt*) encounters the problem that choices that differ in *nAtt* might also differ in difficulty (e.g., Bergert & Nosofsky, 2007; Hunt et al., 2014). To control for difficulty, for each participant *p* we carried out a multiple regression in which the (logarithm of) RTs for each choice type *c* was predicted from *nAtt* and a measure of (inverse) choice difficulty:

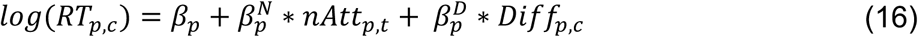

where the (inverse) difficulty measure *(Diff_p_,_c_)* was defined as the absolute value of the log-odds of the probability of a subject choosing alternative A on choice type *c*, computed from the subjective attribute weights derived from the WADD model above,

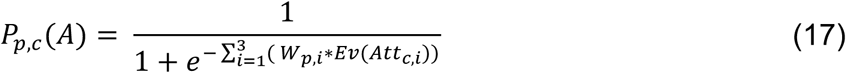

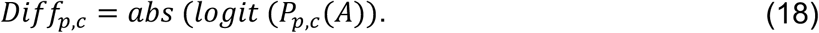

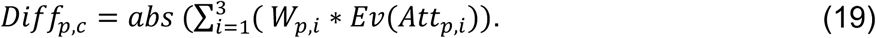

In other words, the greater the evidence in favor of one alternative over the other, the easier the choice. Because it is derived from subjective attribute weights (the *W_p,i_*), *Diff_p,c_* incorporates individual differences in the relative importance of attributes and thus provides a more robust estimate of difficulty as compared to one derived from optimal attribute weights.

Consistent with previous findings (Hunt et al., 2014). RTs indeed decreased with increasing (inverse) difficulty: *β^D^* = −0.101 (SD = 0.144) 0, t(46) = −4.813, p < *10*^−4^. The key result for present purposes is that RTs also increased as the number of discriminating attribute increased, *β^N^* = 0.043 (SD = 0.143), t(46) = 2.063, p < 0.05. This result reflects the fact that participants’ RTs increased by an average of 1.044 s for each additional discriminating attribute. These findings support the notion that choices were evaluated at the level of attributes rather than alternatives.

Note that the preceding analysis included participants in both the full and partial condition. Recall that these two conditions differed in that the former presented 8 distinct types of stimuli whereas the latter presented 20. It is conceivable that the participants were more likely to have formed whole stimulus representations in the full condition in which distinct stimuli were presented more frequently. Therefore, we repeated the analysis in Equation 16 with only the 24 participants in the full condition. For these participants RTs also increased as the number of discriminating attribute increased *β^N^* = 0.074 (SD = 0.142), t(23) = 2.558, p < 0.05, controlling for difficulty, *β^D^* = −0.064 (SD = 0.059), t(23) = −5.330, p < *10*~^−4^. That is, choices were represented at the level of attributes even for participants who were trained on relatively fewer distinct stimuli.

### Model Comparisons

To assess attribute use, choice representation, and the potential presence of recency effect and cue competition quantitatively, we fit each participant’s choice data during the six training blocks and the single test block to each of the six models (Fig. 5A). Model likelihoods were computed from the choice probabilities assigned on every trial. To facilitate model fitting, we used a regularized prior that favored realistic values of inverse temperature (Daw, 2011; Niv et al., 2015). We chose model parameters that minimized the negative log posterior of the data given model parameters. Table 3 shows the average parameter fits for each of the six models. We then computed each participant’s Bayesian Information Criterion (BIC; Schwarz, 1978):

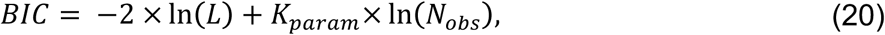

where *L* is the likelihood of the choice probabilities given model and parameter, *K_param_* is the number of parameters in the model, and *N_obs_* is the number of observations (number of trials) for each participant. We then averaged participants’ BICs to compare models. Models that yield a smaller BIC are interpreted as providing a better account of the data.

**Fig. 5.**
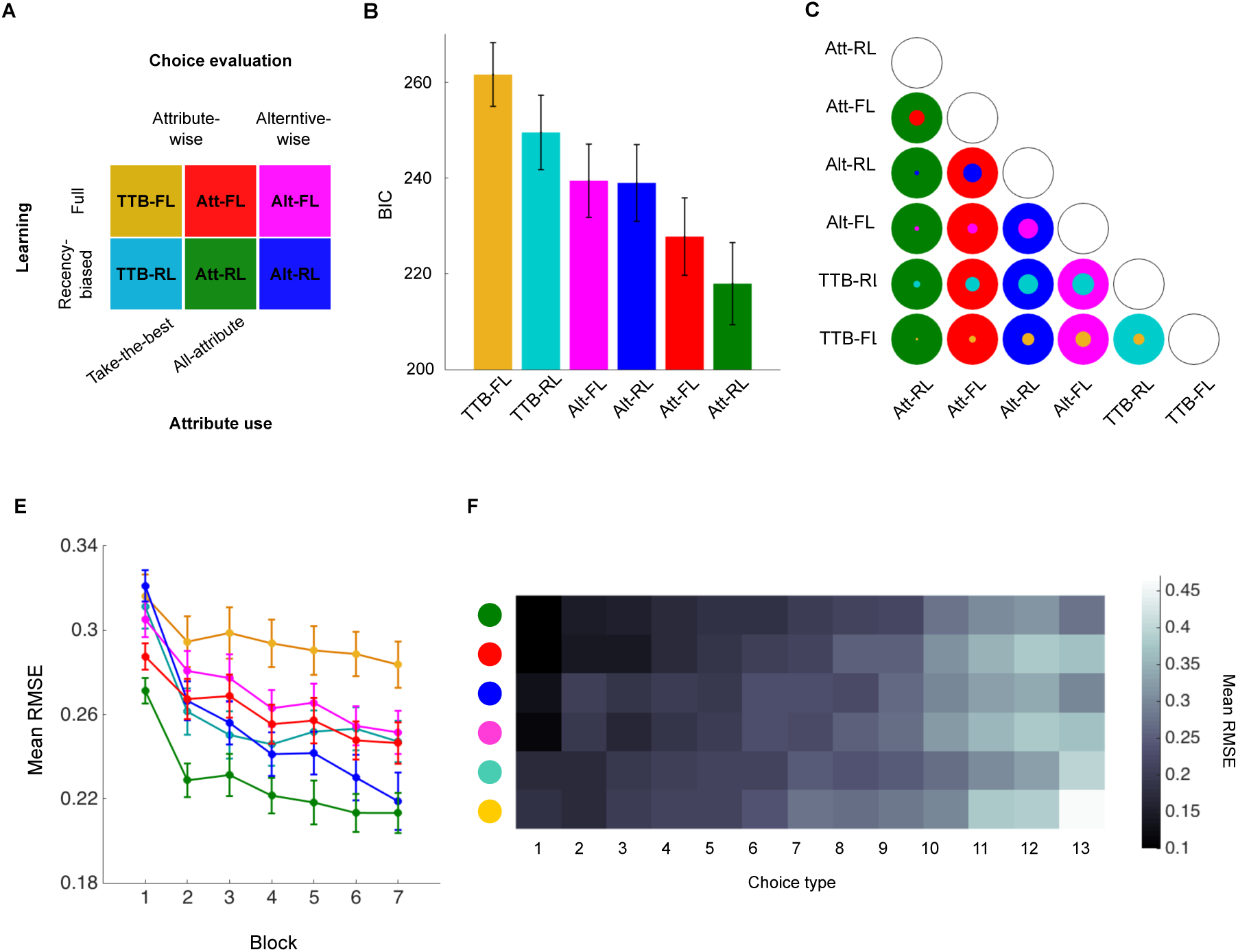
Model comparison. (A) Computational models. Models differ in attribute use, choice representation, and the potential presence of recency effect and cue competition. (B) Model comparison at group level. Mean BICs averaged across participants are displayed. (C) Model comparison at individual level. Each circle illustrates pair-wise comparison between corresponding models. The results are shown in binary colors. The colored area represents the proportion of individual participants that are better fit by each model. (D) Mean root-mean-square error (RMSE) for each model over blocks. (E) Mean RMSE over 13 choice types. The intensity of the color map represents the level of deviation between models’ prediction and participants’ performance. The darker the color, the less the difference (so the better the model).

**Table 3.** Parameter fits to each of the six models.

| **Model** | **Parameter** |  |
| --- | --- | --- |
| | $\alpha$<br>(learning rate) | $\beta$<br>(inverse temperature) |
| TTB-FL | | $0.623 \pm 0.204$ |
| Att-FL | | $0.836 \pm 0.342$ |
| Alt-FL | | $1.064 \pm 0.449$ |
| TTB-RL | $0.166 \pm 0.138$ | $2.126 \pm 1.051$ |
| Att-RL | $0.161 \pm 0.137$ | $2.871 \pm 1.508$ |
| Alt-RL | $0.084 \pm 0.066$ | $3.703 \pm 2.438$ |

Note. Best-fit values were shown as mean ± 1 SD, averaged over individual fits to each participant.

Fig. 5B shows the average BIC values for each of the six models. At the group level, the Att RL model yielded a significantly lower BIC, compared to the second-best model, Att-FL, ΔBIC = 9.839 ± 20.150 (SD), t(46) = −3.347, p < 0.01, which in turn yielded a smaller BIC compared to the third-best model Alt-RL, ΔBIC = 11.192 ± 23.828, t (46) = −3.220, p < 0.01. Although, the BIC differences between models Alt-RL and Alt-FL (ΔBIC = 0.457 ± 21.594, t(46) = −0.145, ns.), and Alt-FL and TTB-RL (ΔBIC = 10.090 ± 41.434, t(46) = −1.670, ns.) were not significant, Alt-RL yielded smaller BIC compared to TTB-RL, ΔBIC = 10.548 ± 33.386, t(46) = −2.166, p < 0.05. Finally, TTB-RL provided significantly better fit compared to TTB-FL, ΔBIC = 12.096 ± 30.375, t(46) = −2.730, p < 0.01. These results indicate that models performed better when choices made use of all attributes, were evaluated attribute-wise, and reflected recency effects and cue competition.

A comparison of the fits of individual participants revealed that models with a lower average BIC also accounted for a greater percentage of individual participants (Fig. 5C). For example, the leading Att-RL model fit better than Alt-FL, Alt-RL, Alt-FL, TTB-RL, and TTB-FL for 70%, 92%, 92%, 87%, and 96% of the participants, respectively; the second-best Att-FL fit better than Alt-RL, Alt-FL, TTB-RL, and TTB-FL for 64%, 80%, 72%, and 87% of participants; and so forth. The leading Att-RL model yielded the best fit for 25 of the 47 participants as compared to 13 for the second-best Att-FL.

Although these analyses identify Att-RL as the best overall account of participants’ choices, it is important to ask whether that advantage obtains over the entire course of the experiment. For example, it is conceivable that Att-RL provided an especially good fit to certain blocks but a poor one to others. To answer this question, we used the maximum likelihood parameters associated with each model and participant to derive the probability of choosing the optimal stimulus for each choice type in each block. We then computed the mean root-mean-square errors (RMSE) between those predictions and participants’ probability of choosing optimally. Fig. 5D presents those RMSEs for each model and block averaged over participants and choice types. This figure reveals that Att-RL is a superior account of participants’ behavior not for a subset of the blocks but rather over the course of the entire experiment.

It is also important to ask whether Att-RL’s advantage obtained for most of the 13 choice types. It is conceivable that Att-RL provided an especially good fit to certain choice types but a poor one to others. To answer this question, Fig. 5E instead averages the RMSEs over blocks (and participants) and so shows how the models compare on the 13 choice types. In fact, Att-RL provided the best or nearly the best account of the large majority of the 13 choice types. Note in particular that it provides the best account of choice type 13, which provides a test of whether participants’ attribute weights are compensatory. The take-the-best models (TTB-FL and TTB-RL) perform poorly on this choice type because they predict that alternative B should be favored. The rational Att-FL model performs poorly because it stipulates compensatory weights and so predicts that alternative A should be favored. The account of choice type 13 provided by Alt-RL is comparable to the one provided by Att-RL, but note that it does much more poorly on many of the other choice types.

The discussion so far describes the relative performance of the six models but not how well those models reproduce participants’ choices. Fig. 6A presents the proportion of optimal choices predicted by each model in each block (colored lines) superimposed on the empirical data (gray bars). Informal inspection of the figure reveals a number of qualitative discrepancies between the performance of the models and the participants. The take-the-best models (TTB-FL and TTB-RL) underestimate participants’ performance in all seven blocks. The rational Att-FL model reaches asymptotic performance too soon, such that learning virtually ceases after Block 4. The alternative-wise models (Alt-FL and Alt-RL) underestimate participants’ performance in Block 1. Overall, the Att-RL provided the best account of participants’ choices over blocks. Nevertheless, note that it also tended to underestimate participants’ early Block 1 performance. The General Discussion will consider potential reasons for this discrepancy.

**Fig. 6.**
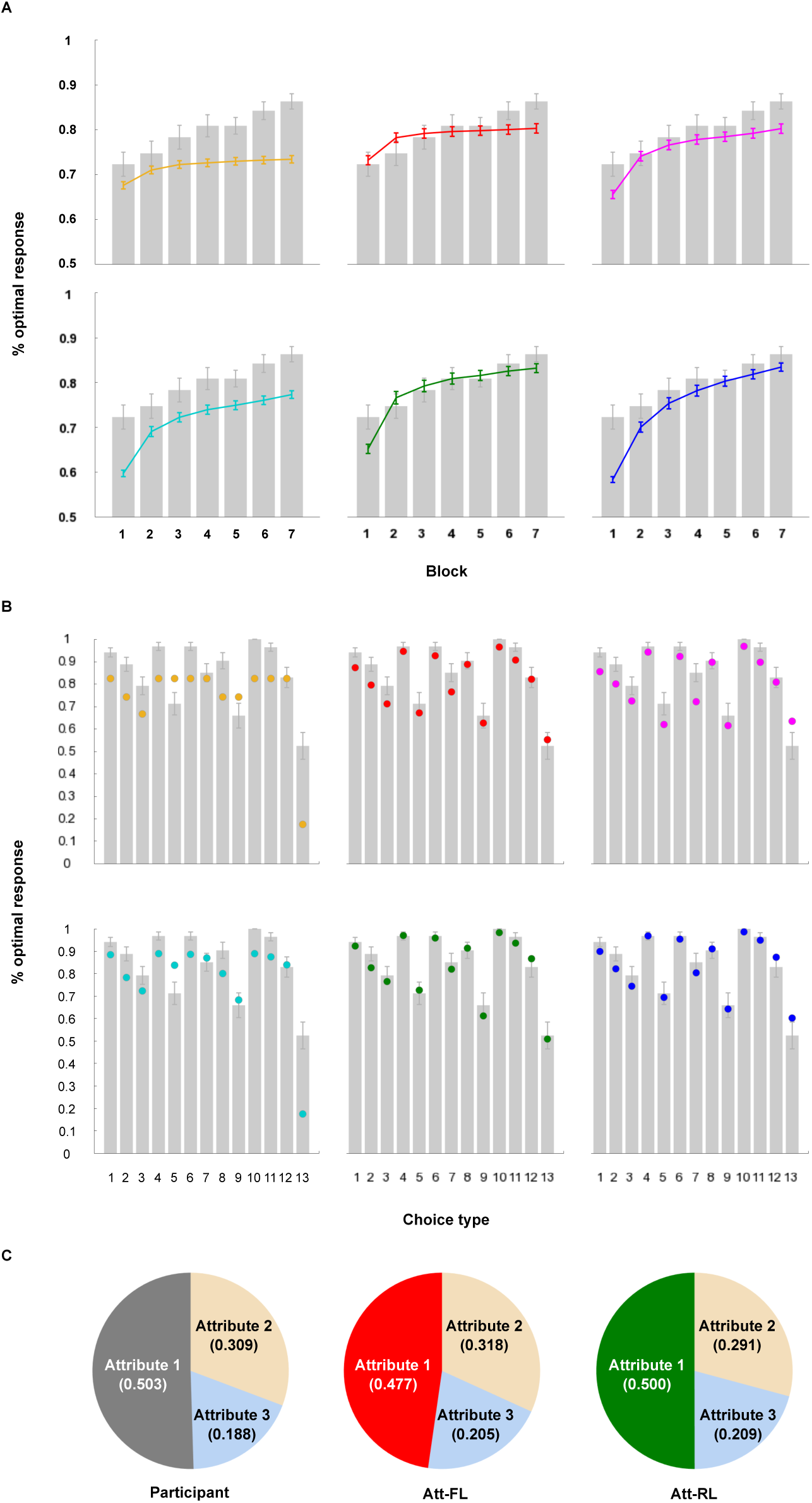
Model performance. (A) Model performance over blocks as compared to that of participants. (B) Model performance over 13 choice types. (C) Normalized attribute weights of participants and those inferred from Att-FL and Att-RL models.

Whereas Fig. 6A considers the models’ fits in each block collapsing over choice types, Fig. 6B presents them for the 13 choice types in the final test block. Note that the data in Fig. 6B is the same as that in Fig. 4B but now model predictions are derived from parameters estimated on the basis of all seven blocks. The figure reveals that the take-the-best models (TTB-FL and TTB-RL) fail to provide a good account of the data. Although the full learning models (Att-FL and Alt-FL) characterize the overall pattern of choices, they substantially underestimate participants’ performance on some choice types (e.g., 1–3, 7, and 11). In contrast, the recency-weighted models (Att-RL and Alt-RL) reproduced participants’ choices quite well. Notably, Att-RL reproduced participants’ 0.5 choice probability on choice type 13, indicating that Att-RL, like the participants, learned to weigh the attributes in a non-compensatory fashion (Fig. 6C). This performance of course can be attributed to the overshadowing of Attribute 3 by the stronger attributes entailed by Att-RL’s error driven learning mechanism. In summary, the overall quantitative measures of fit and the fits to the learning data in Fig. 6A and to choice types in Fig. 6B all identify Att-RL as the best account of participants’ decisions in this experiment.

## Discussion

Taken together, the behavioral results and the computational model fits suggest that participants used multiple attributes when making decisions, that choice alternatives were represented attribute-wise instead of alternative-wise, and that choices reflected RL phenomena such as recency and overshadowing.

One potential caveat regarding these conclusions concerns the fact that participants were instructed at the beginning of the experiment that all three attributes were predictive of reward. One might wonder if learners would have exhibited a qualitatively different strategy (e.g., take the best) in the absence of such instructions. To answer this question, we conducted a follow-up experiment in which participants were instead merely told that “The fact that aliens have different body parts and thus look different from one another will help you identify the rich aliens,” instructions that are neutral regarding the number of predictive attributes. Because the main experiment revealed no differences between the partial and full conditions, only the full condition was tested in the replication. All other aspects of the experiment were unchanged. 12 undergraduate volunteers and 20 paid participants from the New York University community participated the study. Application of the same exclusion criterion used in the main experiment resulted in a total of 20 usable participants. The results were consistent with the main experiment. Participants’ choices reflected the use of multiple attributes (Fig. 7A). Their RTs on three of the four RAT-easy and RAT-hard choice comparisons also implicated the use of multiple attributes (Fig. 7C). We again observed overshadowing in that Attribute 1 was overweighed relative to the other two attributes (Fig. 7B), albeit this difference did not reach significance (a fact likely due to the lower number of subjects). Nonetheless, participants’ at-chance performance on the theoretically important choice type 13 again reflects the non-compensatory nature of their learned attribute weights. Participants’ RTs increased as the number of discriminating attributes increased (*β^N^* = 0.011; SD = 0.024), corroborating our conclusion that choices were evaluated attribute-wise. Finally, we again found that the RL models that incorporate recency and cue competition outperformed the other models (Fig. 7D).

**Fig. 7.**
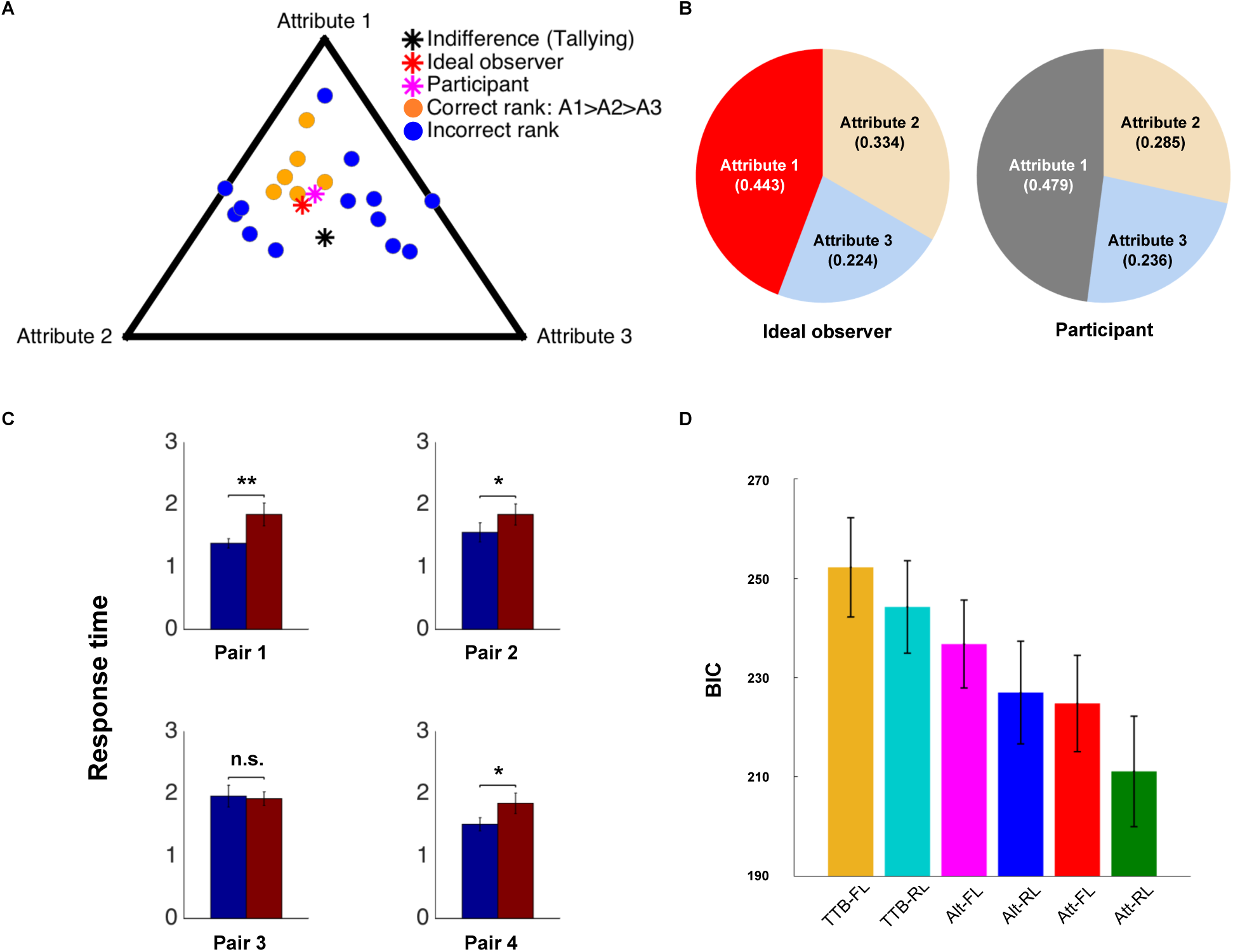
Results from follow-up study. (A) Simplex plot of normalized attribute weights. (B) Average normalized attribute weights inferred from an ideal observer model and that of participants. (C) Response time pairs of choices used to contrast take-the-best and rational models. (D) Model comparison. Mean BIC values are displayed.

## General Discussion

Using a two-alternative choice task with stimuli whose attributes were each predictive of reward, the aim of the present study was to determine what information participants learn to use to make decisions, how they represent that information, and whether those decisions exhibit classic reinforcement learning effects such as recency and cue competition. Analysis of participants’ choices, the time needed to make those choices, and the fitting of computational models together yielded clear answers to all three questions.

Regarding the first question—what information is used—we found that after six blocks of training participants’ were using all three attributes in a weighed additive fashion to make decisions. The question of information use was motivated by the contrast between the rational model, which stipulates that all information is used, and the heuristic take-the-best model, which stipulates that decision makers conserve cognitive resources by choosing solely on the basis of the most discriminating attribute. Although some investigators (Bergert & Nosofsky, 2007; Gigerenzer & Goldstein, 1996; Gigerenzer & Todd, 1999; Rieskamp & Hoffrage, 1999) have concluded that the take-the-best principle characterizes human decision making, universal agreement on this issue is lacking (Bröder, 2000, 2003; Lee & Cummins, 2004; Newell & Shanks, 2003; Newell et al., 2003; Oh et al., 2016; Rieskamp & Hoffrage, 2008; Rieskamp & Otto, 2006). In the present study evidence for the weighted use of multiple attributes came from analyses of participants’ explicit choices as well as their RTs. And, these behavioral indicators were corroborated by model fitting in which versions of the take-the-best model yielded the poorest fit to the data. The present study thus adds to others that conclude that people often do not merely take-the-best when making decisions.

Our evaluation of the take-the-best model as an account of what participants learned was generous in that the attribute weights were assumed to be per-subject free parameters rather than those that should have been induced from the training data. Moreover, because a version of take the best that chooses the best attribute probabilistically is indistinguishable from a linear weighted added model on the basis of choice data alone, we copied the novel RAT-easy and RAT-hard response time analyses introduced by Bergert and Nosofsky (2007). The substantial RT differences between RAT-easy and RAT-hard in our data (Fig. 3D) indicate that our participants’ choices were inconsistent with even the generalized version of the take-the-best model identified by Bergert and Nosofsky.

The poor performance of the take-the-best account may be unsurprising in light of recent observations that this purportedly frugal strategy can be expensive to implement (Bobadilla-Suarez & Love, 2017; Dougherty et al., 2008; Juslin & Persson, 2002). One reason this is so is that although it bases any given decision on the cues of one attribute, it must still learn the optimal weights of all cues (so as to learn which ones are the best, Dougherty et al., 2008). But note that an RL version of take-the-best model that learned attribute weights in a more efficient and so psychologically plausible manner also performed poorly. Our key result is that participants used multiple attributes in a weighted additive fashion, not merely that they used the best attribute as determined by imperfect attribute weights. Take-the-best has also been shown to be inefficient in contexts in which cues appear in unpredictable locations (and thus require search to identify; Bobadilla-Suarez & Love, 2017). But in the present study the stimulus cues (the head, body, and tail of schematic insects) appeared in the same place throughout the experiment. In other words, the manner in which these experiments presented the stimuli was not biased against use of a take-the-best strategy.

Although the current study adds to those that have concluded against the take-the-best strategy, remember that numerous other studies have shown that take the best sometimes characterizes the choices of a subset (e.g., Bröder, 2000, 2003; Lee & Cummins, 2004; Newell & Shanks, 2003; Newell et al., 2003) or the large majority (e.g., Bergert & Nosofsky, 2007) of participants. Factors that have been proposed to promote a take-the-best strategy include task complexity (e.g., the number of cues) (Newell & Shanks, 2003; Newell et al., 2003; Payne, James, & Johnson, 1988), decreasing uncertainty of the environment (Lee & Cummins, 2004; Oh et al., 2016; Shah & Oppenheimer, 2008), use of non-compound versus compound stimuli (in which the cues are features or parts of a single object) (Oh et al., 2016), information or search costs (Bobadilla-Suarez & Love, 2017; Bröder, 2000, 2003; Newell & Shanks, 2003; Rieskamp & Hoffrage, 2008), use of process-tracing procedures (Glöckner & Betsch, 2008; Payne et al., 1988), and time pressure (Bettman, Luce, & Payne, 1998; Oh et al., 2016; Rieskamp & Hoffrage, 2008).

One notable study that concluded in favor of take-the-best is Bergert and Nosofsky (2007), who found that two-thirds of the participants learned to assign 99% of the weight to one attribute. This result is a surprise given the factors cited above as ones that typically promote multiple attribute strategies, namely, the use of compound stimuli and the absence of search costs or time pressure,. We suspect that take-the-best dominated there because many of the subtle cues in their six-dimensional stimuli were perceptually difficult to discriminate and the resulting pressure to conserve cognitive resources led to the use of a heuristic strategy. In contrast, use of TTB is rarer in studies in which cues are easy to discriminate perceptually (e.g., Bobadilla-Suarez & Love, 2017; Oh et al., 2016; and the present one; see Fig. 1).

Perhaps our most striking result is that by the end of learning participants were not only using multiple attributes in a weighted fashion but that they recovered the relative importance of those attributes in a near-optimal way. Besides being remarkable in its own right, this result speaks against the additional heuristic known as tallying, in which decision makers simply count the number of attributes in favor of one alternative without differentially weighing those attributes (Bobadilla-Suarez & Love, 2017; Gigerenzer & Gaissmaier, 2011). Indeed, to the extent that participants chose non-optimally, they did so in a manner in which Attribute 1 was weighed too heavily relative to Attribute 3, with the result that participants failed to recover the compensatory property of the optimal weights (and which we have interpreted as reflecting overshadowing, as discussed below). But apart from this departure from optimal responding, it is clear that participants learned the structure of the task surprisingly well.

Regarding our second question—how choice alternatives are represented—we found evidence that even after six blocks of training participants’ choices were being made on the basis of individual attributes rather than whole alternatives. On one hand, previous neuroimaging research has demonstrated the formation of whole-stimuli brain representations with extensive training (Bayley et al., 2005; Yin & Knowlton, 2006). But more recent findings suggest that attribute values are computed in cortical areas and then passed to the ventromedial prefrontal cortex for value integration (Lim et al., 2013; Suzuki, Cross, & O’Doherty, 2017). Furthermore, behavioral and neural studies have demonstrated the presence of within- and between-attribute comparisons that can be captured by function approximation models (in our case, attribute-wise models) that flexibly make use of task structure (Jones & Cañas, 2010; Niv et al., 2015) but not alternative-wise models (Roe, Busemeyer, & Townsend, 2001; Tsetsos, Usher, & Chater, 2010). In our study as well, we found that participants’ test phase RTs scaled with the number of discriminating attributes, a pattern consistent with attribute-wise models (Dai & Busemeyer, 2014; Hunt et al., 2014). This conclusion was further corroborated by the fact that versions of computational models that assumed attribute-wise representations always outperformed their alternative-wise counterparts.

One variable that may have contributed to this finding is the nature of our stimuli. As compared to our non-compound stimuli in which the three attributes were spatially separated, compound stimuli may be more likely to promote the formation of whole stimulus representations. One study that provides evidence suggestive of this possibility is that of Oh et al. (2016), who found that participants’ choices reflected all four attributes when the cues were presented in an integrated as compared to a spatially segregated manner. But of course this result is also consistent with an attribute-wise strategy in which all attributes receive non-zero weight (as in the present study). An interesting avenue for future research would be to compare compound and non-compound stimuli and apply the RT analysis we have introduced here to determine if the former condition is more likely to yield alternative-wise representations.

It is important to acknowledge that while our RT analysis revealed an effect of the number of attributes, that result implicates the use of attribute-wise representations on some, but not necessarily all, choice trials. In particular, it is conceivable that participants had started, but not yet completed, the process of forming a full set of whole stimulus representations. On this account, had subjects received the additional blocks of training needed for them to memorize whole stimuli then the effect of number of discriminating attributes on test block RTs would have been absent. Indeed, a recent study found that participants initially adopted feature-based (attribute-wise) strategy but gradually switched to an object-based strategy after many training trials (Farashahi et al., 2017). These results were further corroborated by model simulations suggesting a representational transformation over time.

Finally, note the behavioral evidence we present in favor of attribute-wise representations—response times—is indirect. One potential avenue for future research is to combine behavioral experiments with neuroimaging techniques to decode representational patterns in the brain (Cohen et al., 2017; Norman, Polyn, Detre, & Haxby, 2006).

Regarding our third question—the presence of phenomena characteristic of error-driven learning such as recency and cue competition—we found that the reinforcement learning variants of the models consistently provided better accounts of participants’ learning and test data as compared to those that assumed perfect learning of past experiences. This finding is of course consistent with a large body of previous work on reinforcement learning and decision-making (e.g., Bayer & Glimcher, 2005; Daw et al., 2011; Erev & Barron, 2005), and contributes to the understanding of multi-attribute decision-making processes from the perspective of learning, which is essential to choice and adaptive behavior (Shohamy & Daw, 2015).

Perhaps the clearest behavioral marker of error driven learning was the presence of overshadowing that resulted in a weight on the most valid Attribute 1 that was relatively too high and one on the least valid Attribute 3 that was relatively too low. Although participants’ choices were close to those of an ideal learner, overshadowing accounts for the one qualitative departure from optimal responding, namely the fact that chance-level performance in choice type 13 that reflects the non-compensatory property of the learned attribute weights. To our knowledge, this is the first demonstration of the presence of classic cue competition effects in the literature on multi-attribute decision making.

We considered a number of alternative interpretations of this pattern of attribute weights. One is that it reflects our participants’ use of a take-the-best strategy, which, because they determine the large majority of decisions, also predicts that stronger attributes will be overweighed. But of course the multiple tests of take-the-best model we conducted (of both training and test performance, of both choices and response times) all indicated that our participants were not taking the best. A second alternative interpretation of the attribute weights is that they reflected time pressure in which participants had insufficient time to process all attributes; specifically, such pressure may have resulted in Attribute 3 being ignored on a subset of trials, resulting in it having a relatively lower weight (Lee & Cummins, 2004; Oh et al., 2016). On this interpretation, the observed pattern of attribute weights is a decision phenomenon (time pressure) rather than a learning phenomenon (overshadowing). However, recall that we found that on average our participants in less than half the 5 s response window. The apparent absence of time pressure in this experiment bolsters our conclusion that the non-optimal weights learned in this experiment were a consequence of error driven learning.

Other studies have observed attribute weights that might be interpreted as reflecting overshadowing. As mentioned, Oh et al. (2016), who compared how participants learned to choose between four-dimensional stimuli under either high or low time pressure, found that they failed to consider the least valid dimensions when time pressure was high (they “dropped the worse”). Yet, even in the low time pressure condition participants tended to overweigh the most valid attribute and underweigh the rest—that is, their attribute weights can be interpreted as reflecting overshadowing (see their Figures 3 and 7). Nonetheless, note that a response deadline was present even in their low time pressure condition and that that deadline was more stringent (2 s) than ours (5 s) raising the possibility this pattern of attribute weights arose because a subset of their participants ran out of time on a subset of trials. Of course, participants also overweighed stronger attributes in studies that found substantial use of the take the best strategy (e.g., Bröder, 2000, 2003; Bergert and Nosofsky’s, 2007) but in those cases the attributes weights reflect the use of qualitatively different decision strategy rather than overshadowing per se.

Our three key findings—that choices were based on all information, that they were made on the basis of attributes, that they exhibited recency and overshadowing—was summarized by the finding that the model that incorporated all three of these effects, the attribute-wise reinforcement learning model, or Att-RL, provided the best account of the present choice data.

But while we conclude in favor of Att-RL, it is important to remember that this model also diverged from participants’ behavior in one important way, namely, it underestimated participants’ learning in the first training block. One possible reason for this mis-prediction may lie in our assumption that participants’ learning rates remained fixed during the course of the experiment. For example, in the context of probabilistic category learning Kruschke and Johansen (1999) have proposed that categorizers’ learning rates exhibit *annealing* in which learning slows during the course of the experiment. They suggest that annealing occurs in probabilistic categorization because the correct response to the same stimulus changes from trial to trial, which learners interpret as indicating that the error signal is unreliable. If correct, there is no reason to think that this phenomenon doesn’t also arise during probabilistic decision making. For example, if Att-RL were given an extra parameter representing the rate at which the learning rate slowed, it could more accurately account for our participants’ learning curve with an increased inverse temperature parameter (that would allow it to account for Block 1 performance) that didn’t also result in overestimating performance in the later blocks (because of the decreasing learning rate). Indeed, researchers in the field of reinforcement learning have found that participants dynamically adjusted their learning rates to reduce the uncertainty of the environment (Daw et al., 2006; Gershman, 2015).

More importantly, we acknowledge that a full account of the data may require incorporating additional cognitive mechanisms. For example, recent research on reinforcement learning showed that selective attention biased both learning and decision (Farashahi et al. 2017; Leong, Radulescu, Daniel, DeWoskin, & Niv, 2017). Thus it is reasonable to incorporate the operation of selective attention that has played a central role in models of category learning (Kruschke & Erickson, 1994; Kruschke, 1992; Kruschke & Blair, 2000; Love, Medin, & Gureckis, 2004; Nosofsky, 1986; Nosofsky & Palmeri, 1997; Nosofsky, Palmeri, & McKinley, 1994; Rehder & Hoffman, 2005a, 2005b), or even hypothesis testing-like mechanisms in which subjects explicitly abandon use of one subset of dimensions in favor of another (Goodman, Tenenbaum, Feldman, & Griffiths, 2008; Kruschke & Johansen, 1999; Nosofsky et al., 1994). Another possibility for future research is to consider the effect of episodic memory. Although the present work has considered the possibility that whole-stimulus representations form after extensive training, another line of research has suggested that choices are influenced by decision makers sampling over their episodic memory traces of individual trials (Bornstein & Norman, 2017; Duncan & Shohamy, 2016; Murty, FeldmanHall, Hunter, Phelps, & Davachi, 2016; Shadlen & Shohamy, 2016), even when abstract stimuli were presented repeatedly on every trial (Bornstein, Khaw, Shohamy, & Daw, 2017). Theoretical analyses suggest that episodic memory is particularly useful at the early stages of learning when the uncertainty of environment is high (Lengyel & Dayan, 2008; Santoro, Frankland, & Richards, 2016). Within multi-attribute contexts, it has been shown that the retrieval fluency of cues best describes participants’ decisions (Dimov & Link, 2017). Accounts based on sampling from episodic memory challenge traditional RL models that compute a running average of choice values and offer new insights into representational flexibility and behavioral adaptation (Gershman & Daw, 2017).

## Conclusion

The present study assessed multi-attribute decision making in a probabilistic environment. As in many previous studies of choice we asked what attributes are used to make choices, with the finding that participants made weighted use of all attributes in nearly optimal fashion rather than using a heuristic like take-the-best. But inspired by recent research into reinforcement learning, our analyses also established that choice alternatives were represented and evaluated attribute-wise rather than alternative-wise. And, we found that our participants’ learning and decisions were consistent with error-driven reinforcement learning models that predict recency and the cue competition effect known as overshadowing. A computational model that incorporated weighted use of all attributes, attribute-rather than alternative-wise representations, and error-driven learning provided a quite good account of both participants’ choices and their response times.

## Acknowledgments

We thank Jan Drugowitsch for sharing his variational Bayes logistic regression analysis codes online. We thank Aaron Bornstein, Bradley Doll, and Alex Rich for helpful discussion.

## Appendix A

In the partial condition, on some trials an attribute in both alternatives was covered by a piece of leaf (Fig. 1D). Participants were instructed that aliens would appear with one of the three body parts missing occasionally. Specifically, they were told that “Sometimes you can see all three body parts, but other times you can only see two of them, with the third part covered by a piece of leaf.”

The unobservable attributes correspond to one of the non-discriminating attributes in choice types 1 to 9, and instantiations of these choice types in the partial condition were expanded beyond those in the complete condition to incorporate missing attributes. For example, in complete condition choice type 1 (characterized by *Ev*(*Att*_1_) = 1 and *Ev*(*Att*_2_) = *Ev*(*Att*_3_) = 0) could be instantiated in four ways: {aaa, baa}, {aab, bab}, {aba, bba}, or {abb, bbb}, where each set denotes alternatives A and B and each “a” and “b” are the cues that favors A and B, respectively. In the partial condition, there were four additional instantiations: {axa, bxa}, {axb, bxb}, {aax, bax}, or {abx, bbx}, where “x” denotes missing attributes. Thus, in the partial condition choice types 1-3 had eight instantiations and types 4-9 had three (because all attributes discriminate alternatives in types 10-13, so they had the same number instantiations, one, as in the complete condition).

To yield the same number of presentations for each choice type as in the complete condition (Table 1), each instantiation was presented three times for types 1-3 (24 presentations for each type) and four times for types 4-9 (12 presentations for each type). These instantiations were assigned to blocks such that the number of choice type in each block was the same: 4 instances for choice types 1-3, 2 instances for types 4-9 and 3 instances for types 10-13. The reward structure in the partial condition was identical as in the complete condition (Table 1), and an ideal observer would yield the same attribute weights as in the complete condition.

## Footnotes

1 The WADD decision model is traditionally defined in terms of weights on individual cues rather than attributes. However, the structure of our training and test trials were such that the choices predicted by WADD only depend on the *differences* between the weights of the cues on the same dimension. The attribute weights yielded by Eq. 14 can be interpreted as reflecting those differences. Also note that WADD is closely related to what Bergert and Nosofsky (2007) referred to as a *generalized rational model (or gRAT)* in which cue weights are free parameters rather than being assumed to be learned perfectly from the training data.

2 In the take-the-best model with three attribute weights, we assumed that a weight *W* on an attribute entailed weights on the two cues of *W* /2 and −*W* /2. That is, we assumed symmetrical cue weights. The take-the-best model with six cue weights relaxes this restriction. Although this more complex model of course achieved a better fit as compared the model with three attribute weights in absolute terms (average log likelihood of −10.315 vs. −11.129), its fit was worse according to a measure (BIC) that corrects for the number of parameters (45.714 vs. 36.59). There is no pattern of cue weights that results in taking the best yielding an adequate description of participants’ test block choices.

3 It is worth noting that the aggregate fit of the ideal observer model was fairly good. Not only was the average BIC for the ideal model superior to that of WADD (27.298 vs. 29.807), it was the better fitting model for 33 of the 47 participants. Nevertheless, remember that the predictions of the ideal observer model diverged from participants’ choices for theoretically important choice type 13.

